# Advancing the design of the kissing bug kill trap for surveillance of triatomines

**DOI:** 10.1101/2025.05.09.653139

**Authors:** Yuexun Tian, Nadia A. Fernández-Santos, Jose G. Juarez, Henry Esquivel, Andrea M. Moller-Vasquez, María Granados-Presa, Adriana Echeverria, Pamela Pennington, Alejandra Zamora-Jerez, Juan P. Fimbres-Macias, Walter Roachell, Paul A. Lenhart, Theresa Casey, Molly E. Keck, Carolyn L. Hodo, Christopher H. Downs, Sarah C. Sittenauer, Claire Nevins, Sujata Balasubramanian, Carlos Angulo, Carlos Palacios-Cardiel, Ramon Gaxiola-Robles, Tania Zenteno-Savin, Sarah A. Hamer, John H. Borden, Michael G. Banfield, Norma Padilla, Gabriel L. Hamer

## Abstract

Standardized surveillance and control of kissing bugs (Hemiptera: Reduviidae: Triatominae), the insect vectors of the Chagas disease parasite *Trypanosoma cruzi*, which causes Chagas disease, remains difficult. The Kissing Bug Kill Trap consists of solar powered LED lights mounted over a column of black funnels. It operates autonomously to capture, kill and preserve adult triatomines. We conducted experiments from 2022-2024 testing potential ways to improve trap performance, ease of deployment, and minimize cost. Thirteen prototypes evaluated in Texas, Guatemala, and Mexico captured 1,531 triatomines. In 2022-2023 we selected a six-funnel trap suspended from a single support pole with an angle bracket, and with four LED lights and a solar panel mounted above the rain-guard, as a reference trap. In 2023, traps with smaller funnels, blue funnels, and blue lights were inferior to the reference trap based on high by-catch of other arthropods and/or fewer triatomines caught per day. In 2024, traps with more or fewer than six funnels or with LED lights mounted on or below the rain guard did not outperform the reference trap. The experiments added five new triatomine species to the four already known to be caught by the Kissing Bug Kill Trap and revealed differences and similarities in phenology of dispersal flights of *Triatoma gerstaeckeri* over a three-year period in Texas. The reference trap was selected as the pre-commercial prototype, based on its suitability for triatomine surveillance and potential for reducing the risk of *T. cruzi* infection by intercepting dispersing adult triatomines before they reach human habitats.

## Introduction

Triatomines (Hemiptera: Reduviidae: Triatominae) are the insect vectors of *Trypanosoma cruzi*, the agent of Chagas disease throughout the Americas. Surveillance techniques to identify and quantify triatomine populations remain challenging, and the availability of surveillance tools lags behind that for other arthropod vectors such as mosquitoes and ticks. Although pesticides have been used as irritants to flush out triatomines from crevices in homes for surveillance and detection purposes (Reynoso et al. 2017), their high cost poses a challenge for long-term and large-scale monitoring. Various traps have been developed to capture triatomines for different settings. For example, sticky trap can act as a physical barriers, preventing triatomines from entering chicken coops or residences where they may establish populations (Abrahan et al. 2011). Lorenzo et al. 1998 reported a capture of 126 triatomines over five consecutive nights at two chicken-coops using yeast-baited traps, which was significantly more efficient than unbaited traps. In addition to yeast, live mice have also been used as bait, showing high efficiency in nidicolous and domiciliary settings (Noireau et al. 1999, 2002).

While those traps target crawling triatomines, several light-based traps have been developed to capture dispersing adult triatomines. These traps capitalize on the well-established attraction of triatomines to lights (Wisnivesky-Colli et al. 1993, Minoli and Lazzari 2006). In 1964, three black lights hung against a white wall attracted a total of 398 *Triatoma protracta*. Other light trap designs, such as cross panel trap with diode lights, cross-vane trap with UV lights, and white cloth with fluorescent lights, have proven effective in capturing dispersing triatomines under different environmental conditions (Sjogren and Ryckman 1966, Abrahan et al 2011, Updyke and Allan 2018, Wisnivesky-Colli et al. 1993, Rebollar-Tellez et al. 2009). Despite their effectiveness, these traps require considerable time to setup and some require manual monitoring, limiting their uses for routine surveillance or large-scale triatomine control. Therefore, there is a need to develop a standardized trap that is easy to deploy, functions autonomously for long periods, and requires minimal maintenance.

Hamer et al. (2024) described the development of the Kissing Bug Kill Trap for collection, killing, and preservation of adult triatomines. The trap is a modified multiple-funnel trap (Lindgren 1983) forming a vertical silhouette with an attractive LED light source powered by a solar panel mounted on top of the funnel column. The trap functions autonomously by automatically turning on the LEDs each night using a photodiode and captures attracted arthropods in a collection cup attached beneath the lowest funnel, containing propylene glycol as a non-toxic preservative. This trap intercepts flying adult triatomines but not immature stages which lack wings. The trap can be checked at different intervals, depending on operational objectives. Over a three-year development period (2019-2021), Hamer et al. (2024) tested several trap prototypes, demonstrated the capture of four triatomine species, and used the traps to evaluate flight dispersal phenology. We continued trials from 2022 to 2024 to test different lengths of the vertical funnel column and different types and positions of LED light stimuli. As triatomines are reportedly attracted to light in the blue (430-555 nm) wavelengths (Reisenman and Lazzari 2006, Pacheco-Tucuch et al. 2012, Otálora-Luna et al. 2018), we also compared blue and black funnel columns and blue and white LED lights.

## Materials and Methods

Experiments were conducted in Texas, USA, Jutiapa, Guatemala, and Baja California Sur, Mexico (Figure 1, 2). Texas study locations included private properties, public areas, and military training areas in the Lower Rio Grande Valley, San Antonio, Bastrop, and College Station, selected based on documented triatomine presence, landscape variability, and availability of collaborators who could manage weekly trap visits. Experiments were set up as randomized complete blocks, with each location considered a replicate.

**Figure 1.**
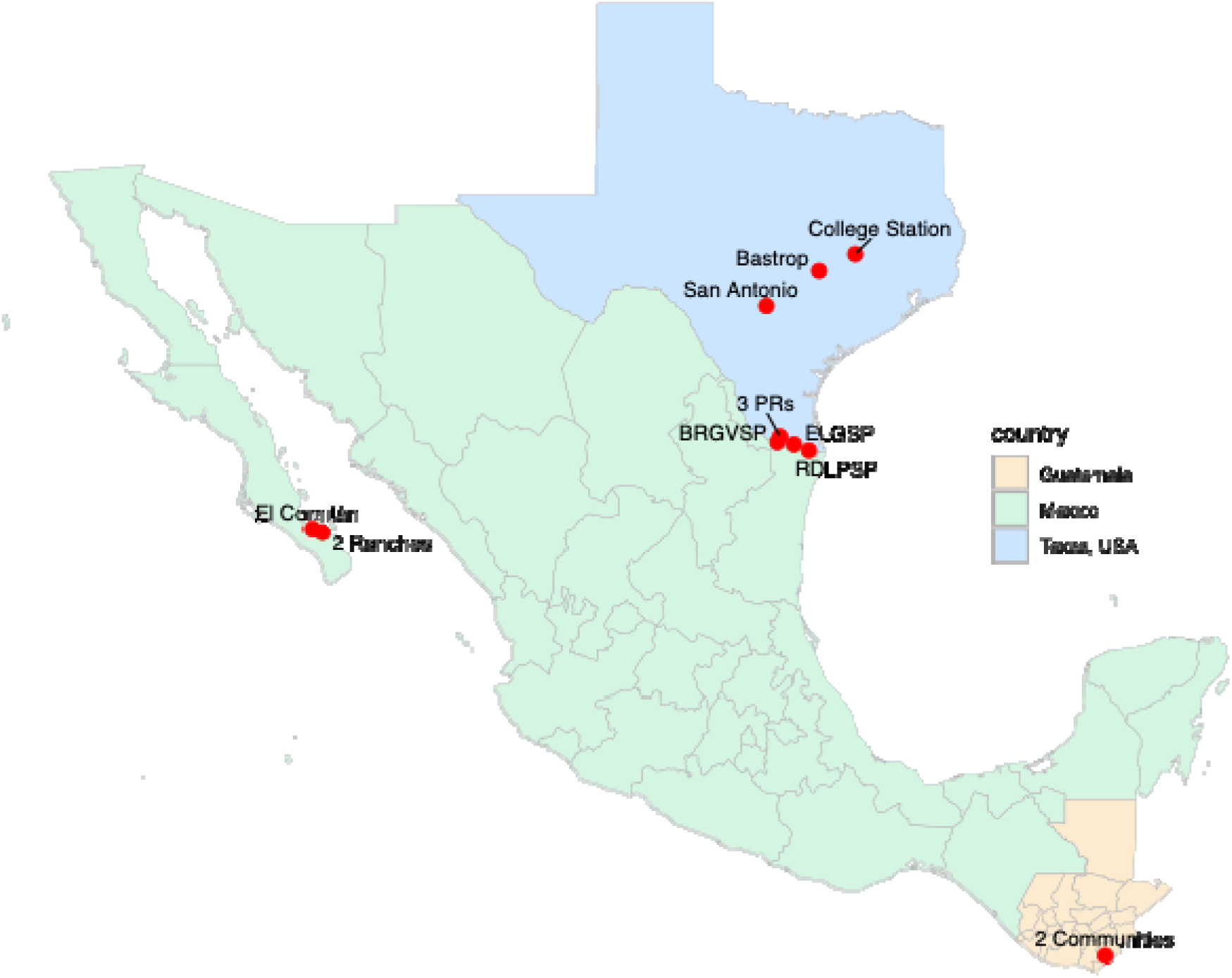
Map of the locations of the traps in Texas, USA, Mexico, and Guatemala during 2022-2024. BRGVSP: Bentsen-Rio Grande Valley State Park; ELGSP: Estero Llano Grande State Park; RDLPSP: Resaca De La Palma State Park; PR: private residence.

**Figure 2.**
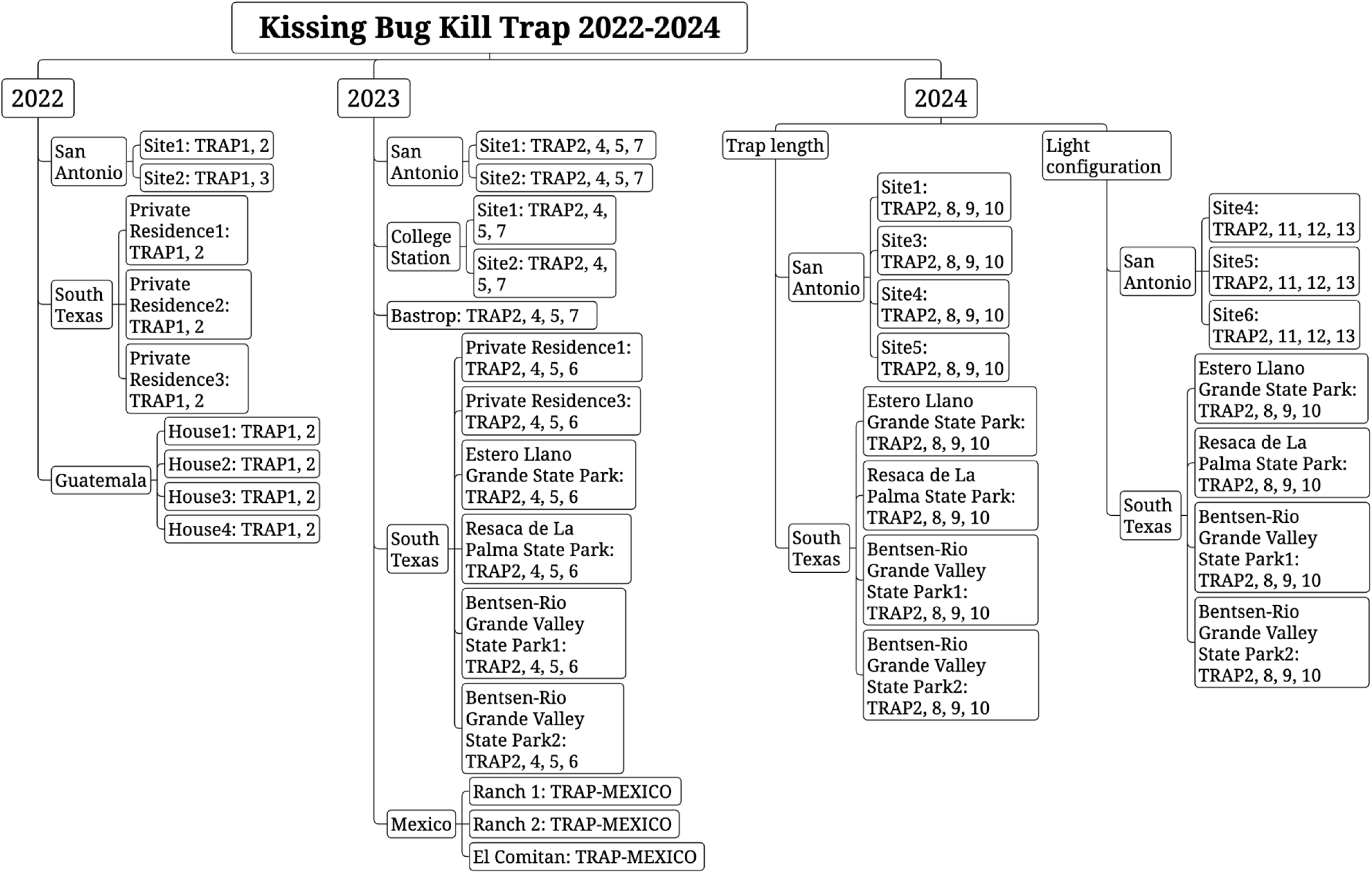
Diagram of the evaluation locations of Kissing Bug Kill Trap in Texas, Guatemala, and Mexico with different trap prototypes (Table 1) deployed from 2022-2024.

The major type of trap used was the Multitrap (Synergy Semiochemicals Corp., Delta, BC, Canada) (Figure 3), a slightly modified Lindgren Funnel Trap, which was designed to be deployed with a semiochemical lure. The Lindgren Funnel Trap was designed to mimic a tree trunk and is especially attractive to forestry pests. It was modified into the Kissing Bug Kill Trap by adding a visual LED lure (Hamer et al. 2024) and was tested with various modifications of the LED light stimulus and length of the funnel column (Table 1). The LED lights in all traps were controlled by a photoswitch providing illumination throughout the scotophase. All collection cups contained propylene glycol (Bluewater Chemgroup, Fort Wayne, IN, USA), a preservative which can be re-used when checking and resetting traps. Traps within each location were spaced at least 50 m apart and were checked every 1-2 weeks. Collected trap contents were transferred to containers with 70% ethanol and transported to a laboratory. Triatomines were separated from by-catch and identified to species (Lent and Wygodzinsky 1979). Other arthropods (by-catch) were identified to order and counted (Supplemental Figure 1).

**Figure 3.**
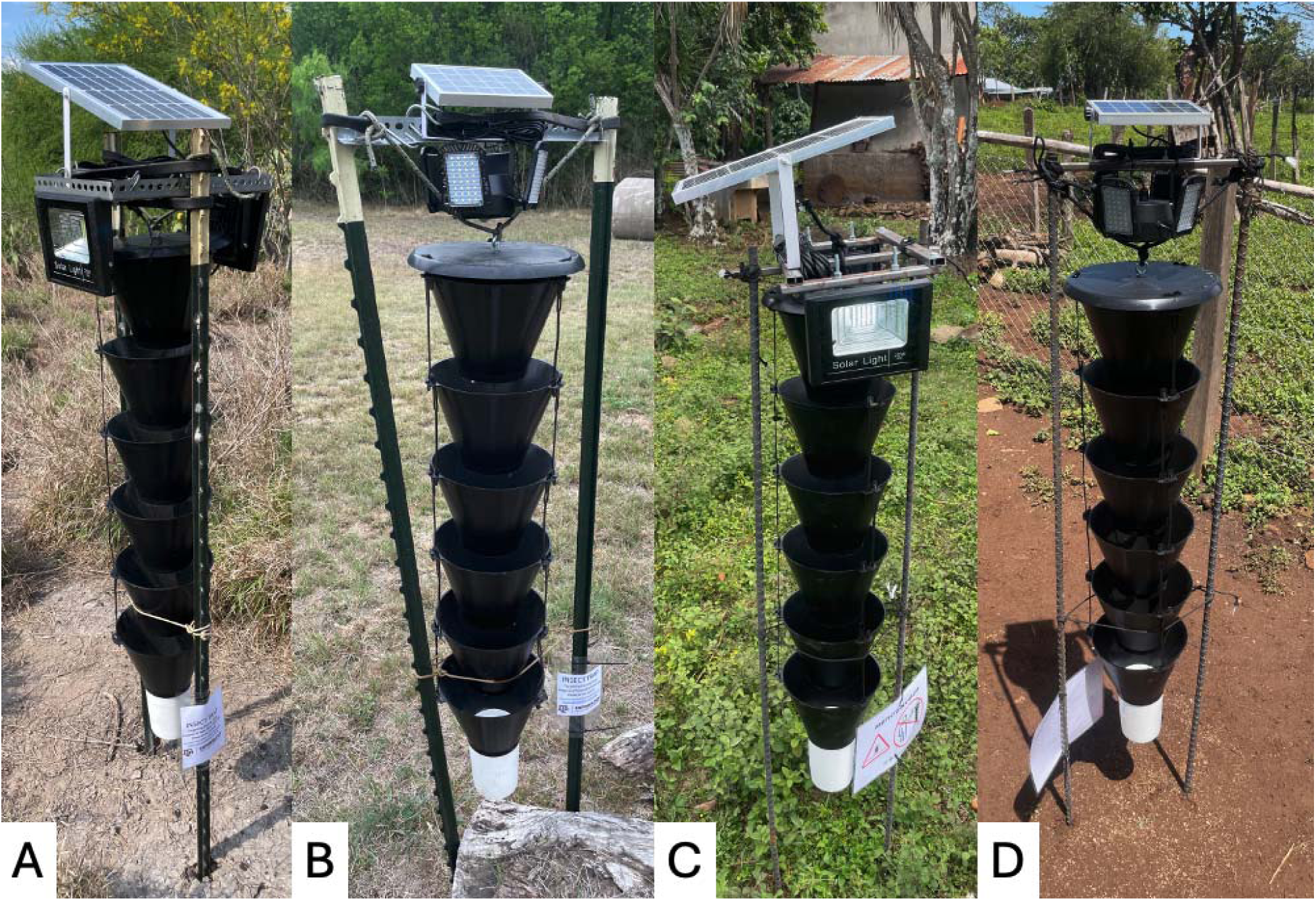
Kissing Bug Kill Trap prototypes evaluated in 2022. A) 6-funnel Multitrap trap with two large light panels deployed in Texas (TRAP1); B) 6-funnel Multitrap trap with 4 small light panels deployed in Texas (TRAP2), C) TRAP1 prototype deployed in Guatemala, D) TRAP2 prototype deployed in Guatemala.

**Table 1.**
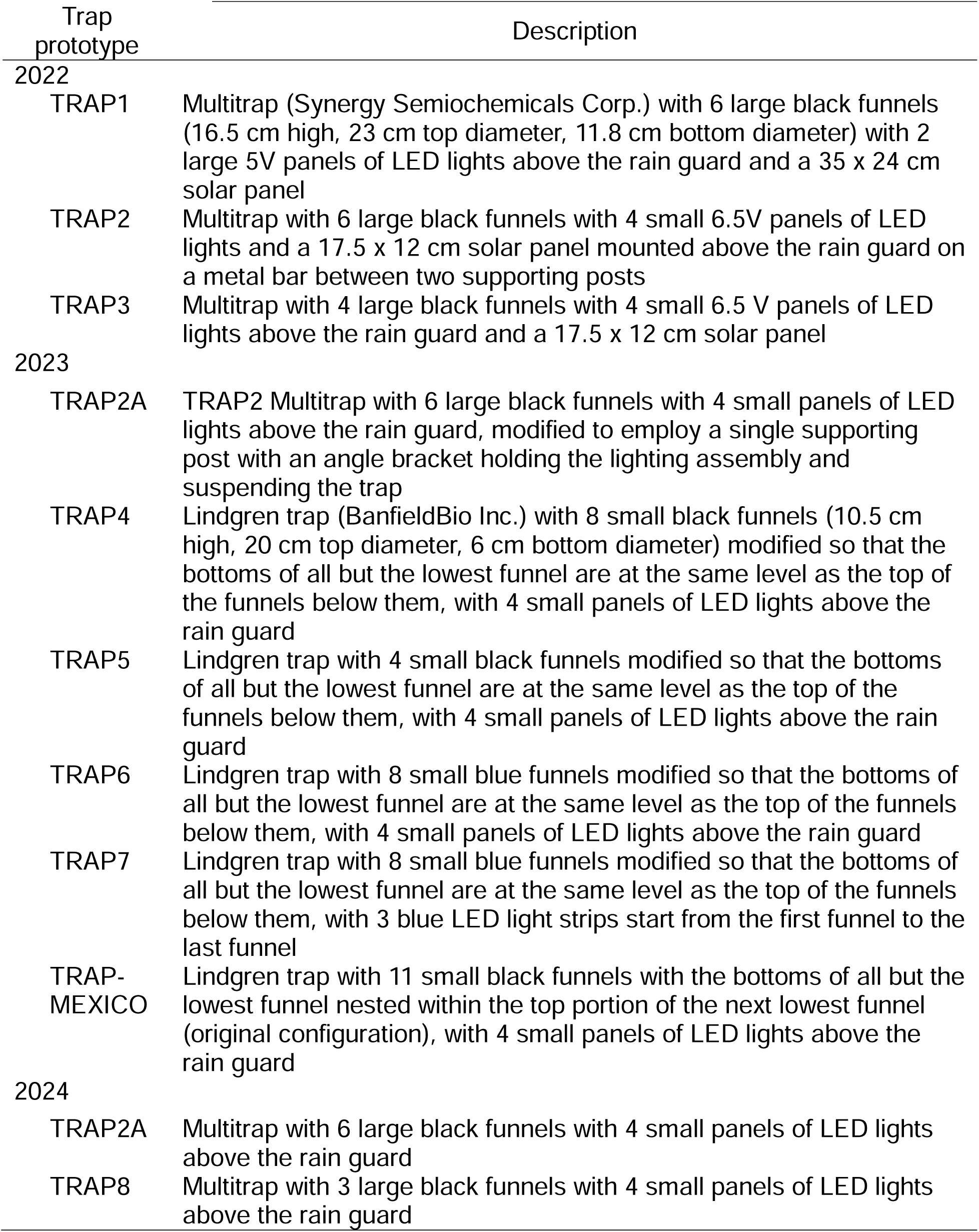
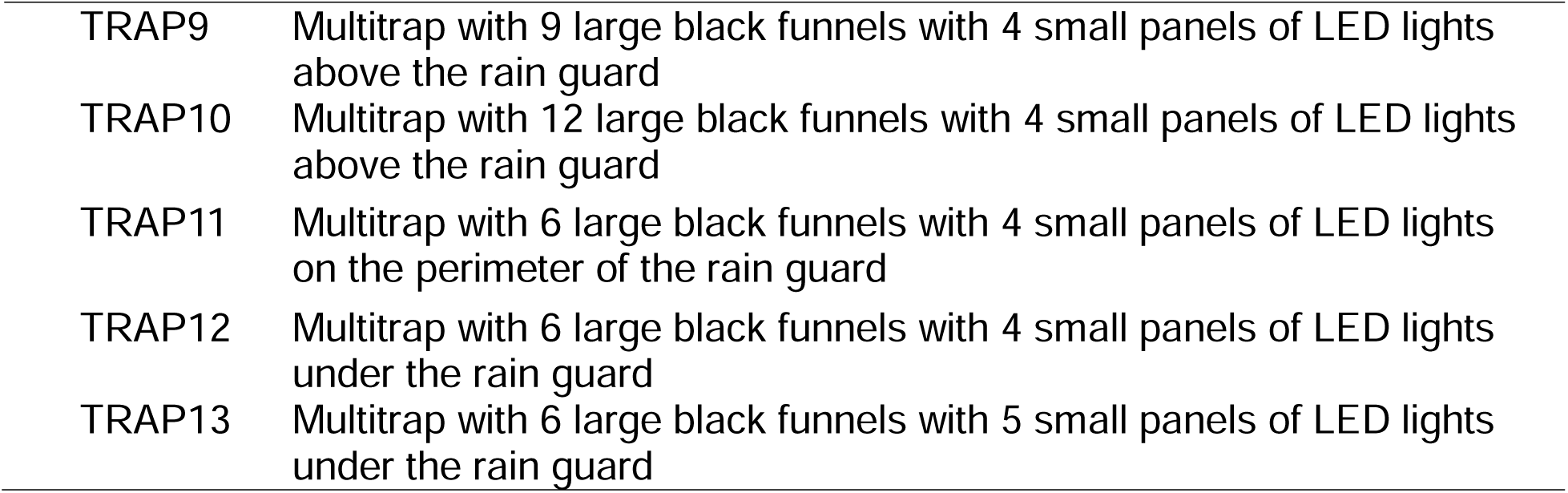
Descriptions of the Kissing Bug Kill Trap in various Multitrap and Lindgren trap modifications tested for capability to capture triatomines in 2022-2024. Modifications are described in trap length, funnel number, size and color, and LED light-solar panel configurations.

| Trap<br>prototype | Description |
| --- | --- |
| 2022 |  |
| TRAP1 | Multitrap (Synergy Semiochemicals Corp.) with 6 large black funnels (16.5 cm high, 23 cm top diameter, 11.8 cm bottom diameter) with 2 large 5V panels of LED lights above the rain guard and a 35 x 24 cm solar panel |
| TRAP2 | Multitrap with 6 large black funnels with 4 small 6.5V panels of LED lights and a 17.5 x 12 cm solar panel mounted above the rain guard on a metal bar between two supporting posts |
| TRAP3 | Multitrap with 4 large black funnels with 4 small 6.5 V panels of LED lights above the rain guard and a 17.5 x 12 cm solar panel |
| 2023 |  |
| TRAP2A | TRAP2 Multitrap with 6 large black funnels with 4 small panels of LED lights above the rain guard, modified to employ a single supporting post with an angle bracket holding the lighting assembly and suspending the trap |
| TRAP4 | Lindgren trap (BanfieldBio Inc.) with 8 small black funnels (10.5 cm high, 20 cm top diameter, 6 cm bottom diameter) modified so that the bottoms of all but the lowest funnel are at the same level as the top of the funnels below them, with 4 small panels of LED lights above the rain guard |
| TRAP5 | Lindgren trap with 4 small black funnels modified so that the bottoms of all but the lowest funnel are at the same level as the top of the funnels below them, with 4 small panels of LED lights above the rain guard |
| TRAP6 | Lindgren trap with 8 small blue funnels modified so that the bottoms of all but the lowest funnel are at the same level as the top of the funnels below them, with 4 small panels of LED lights above the rain guard |
| TRAP7 | Lindgren trap with 8 small blue funnels modified so that the bottoms of all but the lowest funnel are at the same level as the top of the funnels below them, with 3 blue LED light strips start from the first funnel to the last funnel |
| TRAP-MEXICO | Lindgren trap with 11 small black funnels with the bottoms of all but the lowest funnel nested within the top portion of the next lowest funnel (original configuration), with 4 small panels of LED lights above the rain guard |
| 2024 |  |
| TRAP2A | Multitrap with 6 large black funnels with 4 small panels of LED lights above the rain guard |
| TRAP8 | Multitrap with 3 large black funnels with 4 small panels of LED lights above the rain guard |
| TRAP9 | Multitrap with 9 large black funnels with 4 small panels of LED lights above the rain guard |
| TRAP10 | Multitrap with 12 large black funnels with 4 small panels of LED lights above the rain guard |
| TRAP11 | Multitrap with 6 large black funnels with 4 small panels of LED lights on the perimeter of the rain guard |
| TRAP12 | Multitrap with 6 large black funnels with 4 small panels of LED lights under the rain guard |
| TRAP13 | Multitrap with 6 large black funnels with 5 small panels of LED lights under the rain guard |

### 2022: Evaluation of light sources

In 2022, three types of Multitraps were evaluated (Figure 3, Table 1) for the number of light sources and size of the solar panel (TRAP1 *versus* TRAP2 and TRAP3) and length (TRAP1 and TRAP2 *versus* TRAP3). TRAP1 and TRAP2 were deployed at one location in San Antonio and three private residences in south Texas, while TRAP1 and TRAP3 were deployed in another location in San Antonio, resulting in five replicates for TRAP1 (two in San Antonio, three in south Texas), four for TRAP2 (two in San Antonio and two in south Texas, Figure 3), and one for TRAP3. Within each location, suitable trap sites with no shade and good visibility were identified and then the trap type was randomly assigned to them. The traps were installed using two T-posts with different numbers of metal bars supporting the lights above the trap (four bars for large lights and one for small lights) (Hamer et al. 2024). Traps were monitored from April to October 2022 in San Antonio and April 2022 to April 2023 in South Texas.

In Guatemala, eight traps (four TRAP1 and four TRAP2) were deployed on July 7, 2022, in two communities in Comapa, Jutiapa, selected based on previous presence of *T. dimidiata* at low density in households (Bustamante et al. 2014). The traps in Guatemala had the same funnels and lights as Texas but we sourced local materials (1.9-cm rebar and support brackets) to install the traps (Figure 3). In each community, two houses were selected based on field access, location to an open field and safety. In each, we deployed one trap of each type within 10 m of each other. Traps were checked weekly from July 2022 to May 2024. Two manual searches for triatomines using the one-person hour methodology at the households were conducted in May 2023 and May 2024 (Juarez et al. 2018).

### 2023: Color of funnels and light source

TRAP2A was modified from TRAP2 so that the solar panel and LED lights are supported by a single angle bracket and the trap hung from a bracket attached to a single supporting post; this trap was used as a basis for comparison with the Lindgren multiple-funnel trap modified as in Table 1. Because the Lindgren trap has smaller funnels than the Multitrap, eight funnels were needed in TRAP4 to achieve the same column length as TRAP2 with six funnels. TRAP5 was similar to TRAP4, but with only four funnels. TRAP6 was also similar to TRAP4, but the funnels were manufactured from blue plastic containing a pigment that is fluorescent instead of black plastic. TRAP2A, TRAP4, TRAP5, and TRAP6 used the same small 4-panel LED lights. TRAP7 was similar to TRAP6, except that the white LED lights at the top of the trap were replaced with three blue-light LED strips (connected to a 6-volt battery) running vertically down the length of the trap. Suspension of the traps was simplified by using a single T-post with a metal shelving bracket at the top to hold the entire trap (Figure 4).

**Figure 4.**
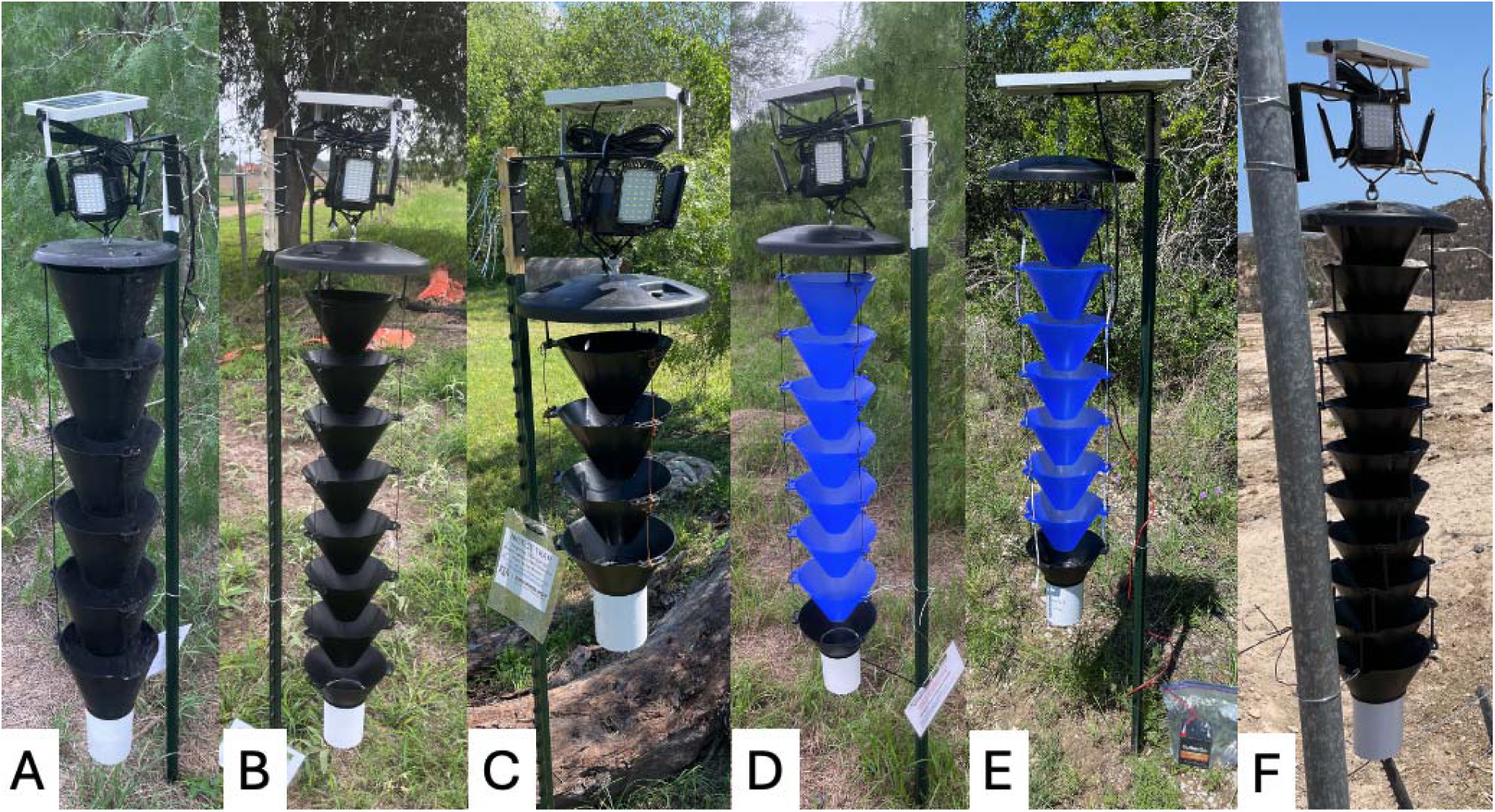
Five Kissing Bug Kill Trap prototypes evaluated in Texas in 2023. A) TRAP2A, Multitrap with 6 funnels and 4-panel lights; B) Lindgren black funnel trap with 8 funnels and 4-panel lights (TRAP4); C) Lindgren black funnel with 4 funnels and 4-panel light (TRAP5); D) Lindgren blue funnel trap with 8 funnels and 4-panel light; E) Lindgren blue funnel trap with 8 funnels and blue LED light strips (TRAP7); F) Lindgren black funnel nested together with 11 funnels and small LED (TRAP-MEXICO).

In central Texas, TRAP2A, TRAP4, TRAP5, and TRAP7, were randomly deployed within two locations in each of San Antonio (eight traps) and College Station (eight traps) and one in Bastrop (four traps), while TRAP2A, TRAP4, TRAP5, and TRAP6 were deployed in each of six locations in south Texas: two private residences (four traps per private residence) and four locations in three state parks [Bentsen-Rio Grande Valley State Park (eight traps), Estero Llano Grande State Park (four traps), Resaca de la Palma State Park (four traps) (Texas Parks and Wildlife State Park Scientific Study Permit No. 26-23)] (Figure 4).

In addition to the five prototypes, another two Lindgren traps of the original design (TRAP-MEXICO, Figure 4, Table 1) were deployed at three locations in Baja California Sur, Mexico. One trap was deployed at Rancho la Huerta on June 20, 2023, taken down on August 22, 2023, and re-deployed at Rancho José Melero on September 13, 2023. One trap was deployed at El Comitán (CIBNOR weather station) on June 22, 2023. Both traps were removed on December 1, 2023, and redeployed at El Comitán from September 1 to November 15, 2024. The traps were checked every 1-4 weeks during these periods.

### 2024: Pre-commercial configurations

Experiments with modified Multitraps in 2024 focused on evaluation of trap length and position of the LEDs. In the first evaluation, TRAP2A with six funnels and small 4-panel LED lights mounted above the rain guard was compared with identical traps with three funnels (TRAP8), nine funnels (TRAP9), and twelve funnels (TRAP10) (Table 1, Figure 5). Traps were randomly deployed in each of the four locations in south Texas (four traps/location) and San Antonio area (four traps/location), resulting in eight replicates for each trap length. In the second evaluation, TRAP2A was compared with traps bearing 4-panel LED lights around the perimeter of the rain guard (TRAP11), 4-panel LED lights below the rain guard (TRAP12), and 5-panel LED lights below the rain guard (TRAP13) (Figure 5). Traps were deployed at three locations in south Texas (three traps/location) and San Antonio area (three traps/location), resulting in six replicates for each light position (Figure 2).

**Figure 5.**
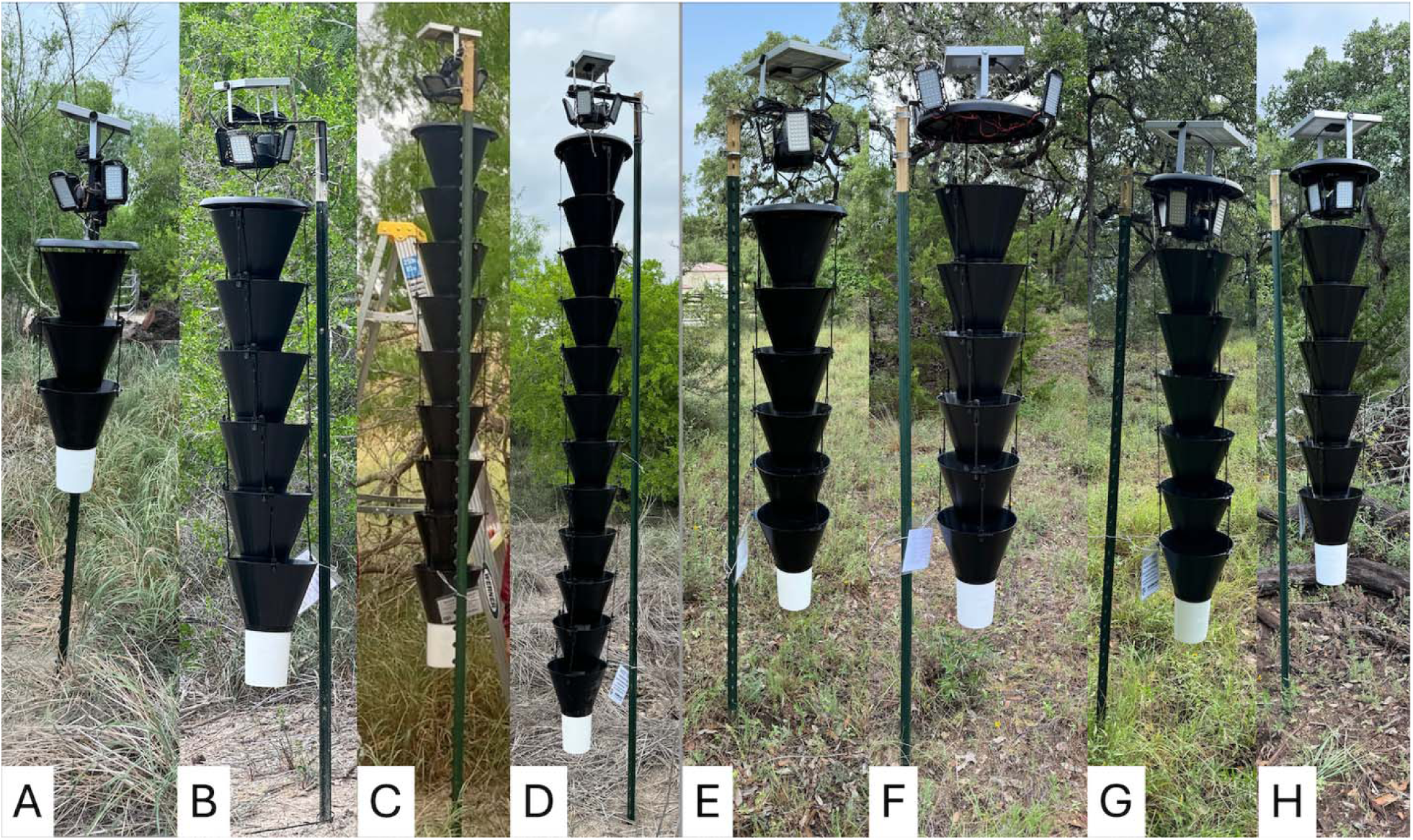
Kissing Bug Kill Traps deployed in Texas in 2024. Group one compares Multitraps with A) TRAP8, 3 funnels, B) TRAP2A. 6 funnels, C) TRAP9, 9 funnels, and D) TRAP10, 12 funnels, and Group 2 compares Multitraps with E) TRAP2A, 4 LED lights above the rainguard, F) TRAP11, 4 LED lights on the perimeter of rainguard, G) TRAP12, 4 LED lights under the rainguard, and H) TRAP13, 5 LED lights under the rainguard.

### Sample processing methods

For traps from the Texas locations with a large number of insects collected, two methods were used to estimate by-catch. The dry weight method was used to process all TRAP7 collections in 2023. After the removal of all triatomines, the remaining trap content was drained, spread evenly in plastic Petri dishes (13.5 cm diameter, 1.8 cm depth), and dried in a fume hood at room temperature for ≥48 h. The desiccated arthropods were then weighed in a new petri dish, and approximately 1/8 of the content was removed, weighed again, and all specimens in the 1/8 content were identified to order. The total number of specimens in each order was estimated based on the equation:

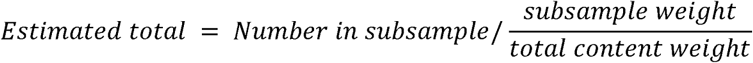

The floating method was used for the 2024 collections to estimate numbers of small arthropods as the small arthropods tend to stick together and were difficult to separate. A round paper of the same 13.5-cm-diameter as a Petri dish was prepared by drawing four lines that equally divided the paper into eight pie pieces (Supplemental Figure 2). This paper was placed under a plastic Petri dish with the trap contents floating inside the Petri dish. The arthropods that were floating in the water were identified and counted individually. The remaining small arthropods that sank to the bottom were estimated by counting and identifying them to order in three randomly selected pie sections. The total number of specimens in each order was estimated based on the equation:

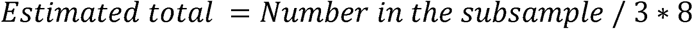

Two triatomines collected from traps deployed in Guatemala were processed with our lab’s established bloodmeal metabarcoding protocol to identify vertebrates fed on prior to capture. These two individuals were also tested for *T. cruzi* using PCR targeting a 166bp region of repetitive microsatellite nuclear DNA. The *T. cruzi* testing and bloodmeal molecular and bioinformatics methods are previously published (Juarez et al. 2025).

### Statistics

Data for each year were analyzed separately. A generalized linear mixed model (GLMM) was used to compare triatomines caught per day and triatomines per 1000 other arthropods among trap prototypes, with the prototypes as variables and the location as random effects. Tweedie distributions were applied because the data were right-skewed and zero-inflated, a common characteristic in ecological count data. Due to loss during transport, some of the triatomines captured in the trap were not identified to species in the lab and were not included in the species-based analyses. The identified triatomine numbers were also analyzed using a GLMM, with a negative binomial distribution, with species and sex as the explanatory variables and years as random effects. A pairwise comparison evaluated the effect of sex on triatomine number within each species. All analyses were conducted using R (R Core Team 2020). Data for which we could not confirm accuracy due to unclear labeling and records were excluded from the analyses.

## Results

During the 3-year evaluation period, the Kissing Bug Kill Trap prototypes captured 1,531 triatomines and 604,432 non-target arthropods (Supplemental Table 1).

### 2022: Evaluation of light sources

In 2022 in Texas, 197 triatomines were captured by the three trap prototypes (five TRAP1, four TRAP2, and one TRAP3) (Table 2). Due to 47 lost specimens during relocation from our field laboratory to main laboratory, only 150 triatomines were identified to species: 49 female and 98 male *T. gerstaeckeri* and 1 female and 2 male *T. sanguisuga*. Mean triatomines caught per day did not differ significantly among the three trap prototypes. However, the mean triatomines per 1,000 other arthropods for TRAP3 (11.288) was significantly higher than for TRAP1 (10.613). Although TRAP2 (6.553), with six funnels and small LED lights, captured fewer triatomines than the prototype with large LED lights and 6 funnels (TRAP1), the difference was not significant. These results were due to the varied number of each trap evaluated (four TRAP2 and one TRAP3), resulting in different variance. Moreover, TRAP2 was light and only required one support metal bar while TRAP1 with large LED lights was heavier and required two metal bars to mount the light and solar panel securely to the stakes.

**Table 2.** Total numbers of triatomines caught and operational days for each trap prototype in Texas in 2022 with means for trap performance and by-catch ratio calculated. A generalized mixed linear model was used to compare trap performance and by-catch ratio with TRAP1 as reference. Asterisk indicates significant difference from TRAP1 (α = 0.05). Dash indicates that the trap was not evaluated at the location. The two San Antionio locations are at the Joint Base San Antonio (JBSA)-Lackland Chapman Training Annex.

|  | TRAP1 | TRAP2 | TRAP3 |
| --- | --- | --- | --- |
| Number of triatomines |  |  |  |
| San Antonio-1 | 14 | 42 | - |
| San Antonio-2 | 65 | - | 17 |
| Private residence-1 (Mission-North) | 10 | 14 | - |
| Private residence-2 (Mission-South) | 20 | 8 | - |
| Private residence-3 (Mission-South2) | 6 | 1 | - |
| Number of operational days |  |  |  |
| San Antonio-1 | 188 | 188 | - |
| San Antonio-2 | 188 | - | 188 |
| Private residence-1 | 357 | 358 | - |
| Private residence-2 | 71 | 71 | - |
| Private residence-3 | 279 | 279 | - |
| Mean triatomines per day | 0.150 | 0.095 | 0.090 |
| Mean triatomines per 1,000 other arthropods | 10.613 | 6.553 | 11.288* |

In Comapa, Guatemala, 13 triatomines, all *T. dimidiata*, were captured from the eight traps during 2022-2024 (Table 3). The traditional manual searches of four homes found no triatomines. Two *T. dimidiata* (collected on October 1 and November 27 of 2022) were processed for bloodmeal metabarcoding. Only one (female) yielded a PCR amplicon of the vertebrate 12S rRNA gene followed by amplicon deep sequencing, which matched *Canis lupus familiaris* (domestic dog) and *Gallus gallus* (chicken). This same female (TD22CompaKB1) was also infected with *T. cruzi* (untypable).

**Table 3.** Total number of triatomines caught and operational days of each trap prototype in Guatemala from 2022-2024 with mean trap performance and by-catch ratio calculated. A generalized mixed linear model disclosed no significant differences in trap performance and by-catch ratio with TRAP1 as reference.

|  | TRAP1 | TRAP2 |
| --- | --- | --- |
| Number of triatomines |  |  |
| House 1 | 1 | 2 |
| House 2 | 1 | 0 |
| House 3 | 5 | 1 |
| House 4 | 3 | 0 |
| Number of operational days |  |  |
| House 1 | 660 | 667 |
| House 2 | 653 | 660 |
| House 3 | 660 | 667 |
| House 4 | 667 | 667 |
| Trap performance (Average triatomine per day) | 0.004 | 0.001 |
| By-catch ratio (Average triatomine per 1,000 other arthropods) | 0.181 | 0.066 |

### 2023: Color of funnels and light source

In 2023 in Texas, 44 traps of 5 prototypes caught 492 triatomines with 452 of them identified to species, including 77 female and 346 male *T. gerstaeckeri*, 7 female and 9 male *Paratriatoma lecticularia*, 3 female and 4 male *Hospesneotomae neotomae* [formally known as *Triatoma neotomae* (de Paiva et al. 2025)], and 3 female and 3 male *T. sanguisuga* (Table 4)). TRAP4 (with 8 small black funnels) and TRAP5 (with 4 small black funnels) caught significantly fewer triatomines than TRAP2A (with 6 large black funnel). TRAP7 (with blue funnels and blue light) had significantly fewer (p<0.01) triatomines per other arthropods than TRAP2A.

**Table 4.** Total number of triatomines caught and operational days of each trap prototype in Texas in 2023 with mean trap performance and by-catch ratio calculated. A generalized mixed linear model was used to compare trap performance and by-catch ratio with TRAP2A trap as reference. Asterisk indicates significant difference from TRAP2A (*P* = 0.05). The two San Antionio locations are at the Joint Base San Antonio (JBSA)-Lackland Chapman Training Annex.

|  | TRAP2A | TRAP4 | TRAP5 | TRAP6 | TRAP7 |
| --- | --- | --- | --- | --- | --- |
| Number of triatomine |  |  |  |  |  |
| San Antonio-1 | 7 | 4 | 5 | - | 3 |
| San Antonio-2 | 4 | 4 | 6 | - | 6 |
| College Station-1 | 0 | 0 | 0 | - | 0 |
| College Station-2 | 0 | 0 | 0 | - | 0 |
| Bastrop | 3 | 3 | 1 | - | 1 |
| Estero Llano Grande State Park | 96 | 10 | 9 | 21 | - |
| Resaca De La Palma State Park | 53 | 45 | 6 | 13 | - |
| Bentsen-Rio Grande Valley State Park -1 | 0 | 6 | 2 | 20 | - |
| Bentsen-Rio Grande Valley State Park -2 | 85 | 13 | 8 | 12 | - |
| Private residence-1 | 3 | 14 | 9 | 19 | - |
| Private residence-4 | 1 | 0 | 0 | 0 | - |
| Number of operational days |  |  |  |  |  |
| San Antonio-1 | 63 | 62 | 49 | - | 56 |
| San Antonio-2 | 71 | 61 | 67 | - | 42 |
| College Station-1 | 95 | 95 | 95 | - | 59 |
| College Station-2 | 108 | 101 | 99 | - | 81 |
| Bastrop | 129 | 129 | 129 | - | 129 |
| Estero Llano Grande State Park | 380 | 367 | 317 | 372 | - |
| Resaca De La Palma State Park | 373 | 373 | 334 | 373 | - |
| Bentsen-Rio Grande Valley State Park -1 | 368 | 368 | 336 | 368 | - |
| Bentsen-Rio Grande Valley State Park -2 | 372 | 372 | 335 | 372 | - |
| Private residence-1 | 368 | 359 | 332 | 368 | - |
| Private residence-4 | 369 | 369 | 333 | 349 | - |
| Trap performance (Average triatomine per day) | 0.075 | 0.035* | 0.027* | 0.038 | 0.041 |
| By-catch ratio (Average triatomine per 1,000 other arthropods) | 5.052 | 3.445 | 3.9102 | 3.723 | 0.072* |

Four triatomines were captured from the two traps in Mexico. Two *Dipetalogaster maxima* (one female and one male) were captured at Rancho la Huerta on July 21, 2023. One female and one male *Hospesneotomae peninsular* [formerly known as *Triatoma peninsular* (de Paiva et al. 2025)] were captured at El Comitán (CIBNOR weather station) on October 18, 2024.

### 2024: Pre-commercial configurations

In 2024 in Texas, 690 triatomines were collected, 333 in the 32 traps with different numbers of funnels and 357 in the 24 traps with different light configurations (Tables 5,6). The 626 triatomines identified to species included 175 female and 411 male *T. gerstaeckeri*, 2 female and 4 male *T. indictiva*, 8 female and 8 male *P. lecticulari*, 3 female and 3 male *H. neotomae,* and 2 female and 5 male *T. sanguisuga,* with 5 specimens too damaged to be identified.

Traps with 3, 9, or 12 funnels did not catch a significantly different number of triatomines than TRAP2A (Table 5). TRAP2A caught significantly more triatomines per day and significantly more triatomines per 1,000 other arthropods than traps with the LED lights on the perimeter of (TRAP11) or below (TRAP12, TRAP13) the rain guard (Table 6). However, increasing the number of LED lights from four to five in TRAP13 raised the number of triatomines caught per day to a level not significantly different from that in TRAP2A.

**Table 5.** Total number of triatomines caught and operational days of trap prototypes with different numbers of funnels in Texas in 2024 with mean trap performance and by-catch ratio calculated. A generalized mixed linear model disclosed no significant differences in trap performance and by-catch ratio with the TRAP2A trap as reference. San Antonio-1 and San Antonio-3 are at the Joint Base San Antonio (JBSA)-Lackland Chapman Training Annex.

|  | TRAP2A | TRAP8 | TRAP9 | TRAP10 |
| --- | --- | --- | --- | --- |
| Number of triatomine |  |  |  |  |
| San Antonio-1 | 1 | 0 | 8 | 3 |
| San Antonio-3 | 7 | 1 | 0 | 3 |
| San Antonio-4 | 0 | 1 | 3 | 7 |
| San Antonio-5 | 1 | 1 | 4 | 1 |
| Resaca De La Palma State Park | 4 | 4 | 25 | 13 |
| Bentsen-Rio Grande Valley State Park -1 | 8 | 42 | 11 | 17 |
| Bentsen-Rio Grande Valley State Park -2 | 56 | 2 | 24 | 16 |
| Estero Llano Grande State Park | 12 | 9 | 23 | 26 |
| Number of operational days |  |  |  |  |
| San Antonio-1 | 113 | 114 | 113 | 113 |
| San Antonio-3 | 107 | 113 | 113 | 110 |
| San Antonio-4 | 113 | 113 | 113 | 113 |
| San Antonio-5 | 114 | 114 | 114 | 114 |
| Resaca De La Palma State Park | 215 | 215 | 215 | 191 |
| Bentsen-Rio Grande Valley State Park -1 | 214 | 214 | 214 | 209 |
| Bentsen-Rio Grande Valley State Park -2 | 214 | 214 | 214 | 214 |
| Estero Llano Grande State Park | 214 | 214 | 214 | 214 |
| Trap performance (Average triatomine per day) | 0.057 | 0.037 | 0.065 | 0.059 |
| By-catch ratio<br>(Average triatomine per 1,000 other arthropods) | 0.431 | 0.311 | 0.731 | 0.566 |

**Table 6.** Total number of triatomines caught and operational days of trap prototypes with different light configurations in Texas in 2024 with mean trap performance and by-catch ratio calculated. A generalized mixed linear model was used to compare trap performance and by-catch ratio with the TRAP2A trap as reference. Asterisk indicates significant difference from TRAP2A (*P* = 0.05).

|  | TRAP2A | TRAP11 | TRAP12 | TRAP13 |
| --- | --- | --- | --- | --- |
| Number of triatomine |  |  |  |  |
| San Antonio-4 | 4 | 2 | 2 | 4 |
| San Antonio-5 | 5 | 1 | 4 | 2 |
| San Antonio-6 | 32 | 16 | 24 | 34 |
| Resaca De La Palma State Park | 76 | 4 | 20 | 30 |
| Bentsen-Rio Grande Valley State Park | 10 | 3 | 3 | 11 |
| Estero Llano Grande State Park | 22 | 6 | 18 | 24 |
| Number of operational days |  |  |  |  |
| San Antonio-4 | 69 | 69 | 69 | 69 |
| San Antonio-5 | 69 | 69 | 69 | 69 |
| San Antonio-6 | 72 | 69 | 72 | 69 |
| Resaca De La Palma State Park | 171 | 171 | 171 | 171 |
| Bentsen-Rio Grande Valley State Park | 164 | 164 | 164 | 164 |
| Estero Llano Grande State Park | 164 | 164 | 164 | 164 |
| Trap performance (Average triatomine per day) | 0.202 | 0.059* | 0.111* | 0.161 |
| By-catch ratio<br>(Average triatomine per 1,000 other arthropods) | 0.807 | 0.284* | 0.282* | 0.359* |

### Triatomine species and sex ratios

*Triatoma gerstaeckeri* was caught in significantly higher numbers (z = 5.311, *P* < 0.001) compared to all other species (Figure 6), with 94.1% (n = 1,156) of all captures. There was no interaction between sex and triatomine species (*P* >0.1). Pairwise comparisons revealed 2.8x more *T. gerstaeckeri* males than females captured (*P* <0.01) but there was no difference in catch by sex for *T. indictiva* (*P* = 0.30), *P. lecticularia* (*P* = 0.51), *H. neotomae* (*P* = 0.79), and *T. sanguisuga* (*P* = 0.48).

**Figure 6.**
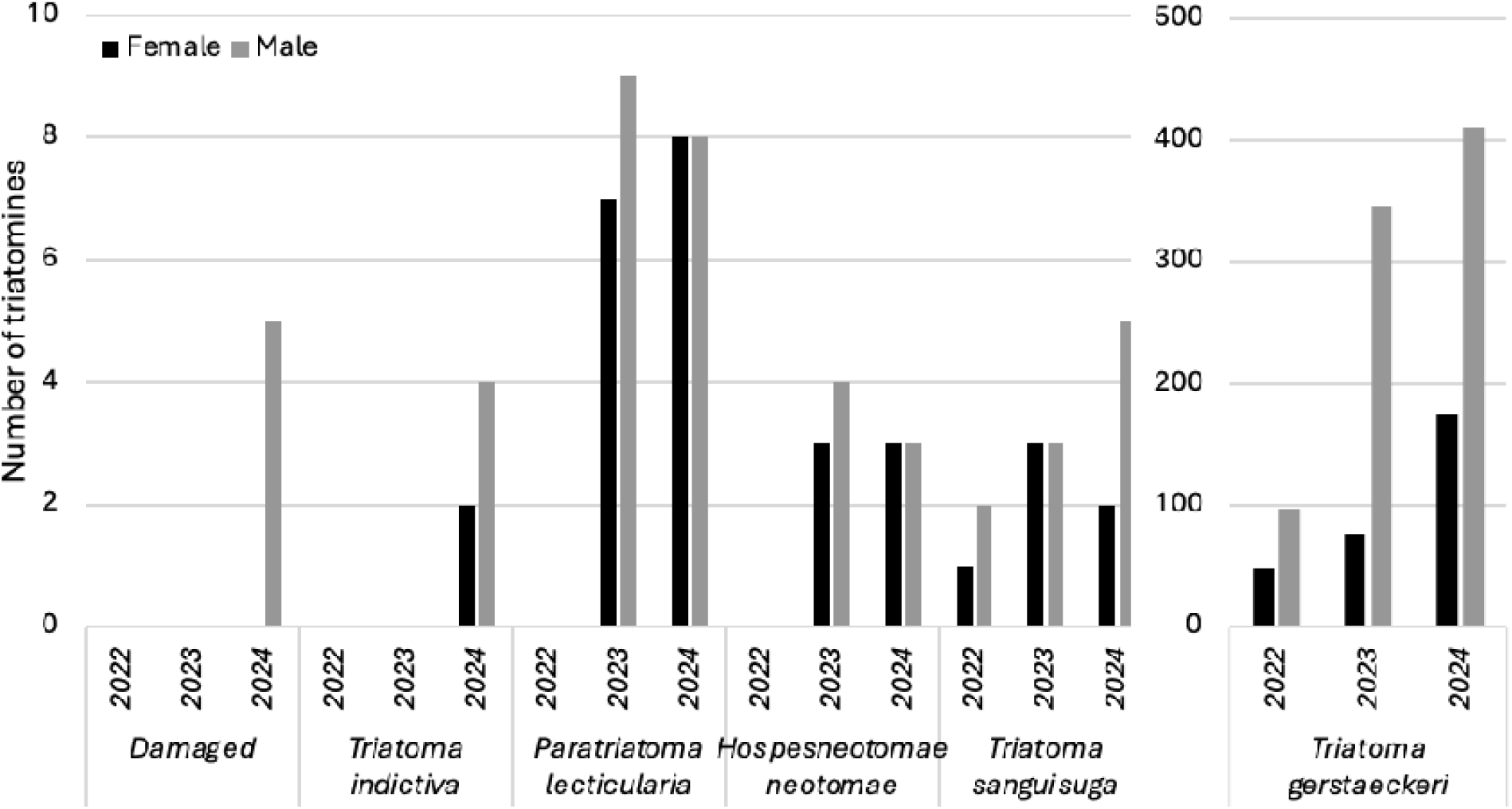
Number of each triatomine species and sex collected from the traps from 2022-2024.

### Phenology

Adult flight dispersal phenology for *T. gerstaeckeri* plotted as the numbers of bugs caught per day per trap among the traps for each week of sampling differed between years and regions (Figure 7). Dispersal flights in 2022 had begun in both south Texas and San Antonio when traps were deployed in April, and dispersal activity in both places diminished by the first week of August. The 2023 and 2024 dispersal flights in South Texas both began in the third week of March. The 2023 flight ended in the third week of September, while the 2024 flight extended into October. In both years flight in San Antonio began several weeks later and ended several weeks earlier than in South Texas.

**Figure 7.**
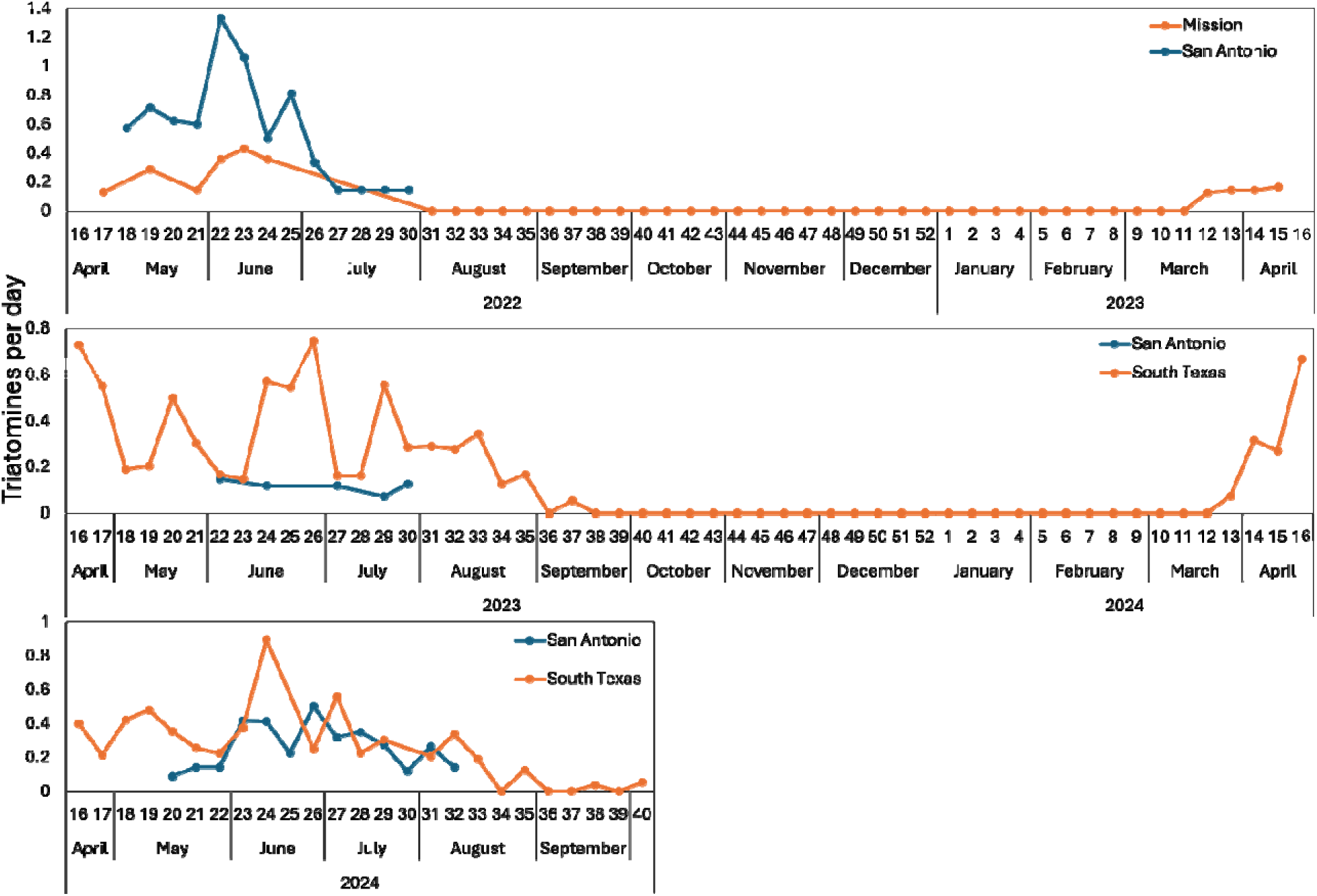
Weekly phenology of *Triatoma gerstaeckeri* catch per day per trap in 2022-2024 at south Texas and San Antonio. The trap data for all south Texas locations and all San Antonio locations are averaged.

Other triatomine species in Texas were captured in smaller numbers than *T. gerstaeckeri* (Supplemental Table 2). *Triatoma sanguisuga* were captured from May to August, *H. neotomae* were captured from April to October, *T. indictiva* were captured in July and August, and *P. lecticularia* were captured from April to September. Thirteen *T. dimidiata* were captured throughout the year in Guatemala, three during the rainy season from May to October and 10 during the dry season from November to April.

### Trap by-catch and failures

The by-catch, measured as triatomine per 1,000 other arthropods, ranged from 0.04 for TRAP7 (blue funnels with blue LED light strips) to 11.29 for TRAP3 (4 black funnels and 4 LED lights) for the 15 trap prototypes that received the full processing of total arthropods (Supplemental Table 1). While the TRAP7 blue funnel and blue light trap was only operated for one season in Texas, this trap captured 261,495 individual arthropods, which is 43.3% of all the by-catch of all 15 prototypes for the entire three years study. The longitudinal pattern in by-catch of other arthropods showed peaks in in early June and late August in South Texas, 2022 and in June in San Antonio in 2023 and 2024 (Figure 7).

Over the three-year evaluation period of 15 traps, occasional malfunctions were observed and promptly addressed. For example, connections on one 12-funnel trap in 2024 broke and the trap fell, which were repaired to restore functionality. Similarly, malfunctioning lights were either fixed or replaced as soon as the issue was detected. In a single instance in 2024, the solar panel light from one trap was stolen from one site in San Antonio (Lackland Chapman Training Annex); we replaced the missing light with a backup unit to maintain continuous operation. For all trap failures, any collection data obtained from the affected trap during the malfunction period were discarded and were not included in analyses.

## Discussion

Over a period of three years (2022-2024), we evaluated 13 Kissing Bug Kill Trap prototypes in three countries which captured 1,531 triatomines of eight species. In 2022, TRAP2 with six black funnels, two support posts and four small LED light panels mounted above the rain guard on a bar suspended between the support posts was comparable to the same trap with two large light panels (TRAP1) in triatomines caught per day but was inferior in triatomines caught per 1,000 other arthropods. This inferiority was not found in later experiments. Moreover, the LED lights and solar panel of TRAP2 were lighter than the larger LED lights and solar panel of TRAP1, which required a cumbersome square frame mounted to the two support posts. Because TRAP2 was lighter, cheaper, and easier to install than TRAP1, it was modified by employing a single supporting post with an angle bracket holding the lighting assembly and suspending the trap (called TRAP2A). TRAP2A was selected as a reference treatment for experiments in 2023 and 2024. These experiments upheld its superiority.

In 2023, TRAP2A outperformed all other traps tested regardless of color or length of the funnel column, and in 2024 it outperformed other Multitrap prototypes with either different numbers of funnels or lower LED light positions. Lindgren traps were designed to catch flying beetles (Lindgren 1983) and the modifications incorporated into Multitraps improved their efficacy for this use (Miller et al. 2013). Unlike the original Lindgren trap [designated as TRAP-MEXICO (Figure 4F)] which had nested funnels, other versions of the Lindgren trap in our experiments had the funnels configured so that the bottom of one funnel was at the same level as the top of the funnel below it (Figure 4B-E), just as the funnels in TRAP2 and TRAP2A were configured. Therefore, other features of TRAP2 and TRAP2A would have been responsible for their superior performance. These features include wider funnels and apertures, greater funnel height and a steeper slope.

Despite evidence that triatomines are attracted to blue light (Reisenman and Lazzari 2006, Pacheco-Tucuch et al. 2012, Otálora-Luna et al. 2018), blue Lindgren traps with white LED lights (TRAP6) or blue LED strips (TRAP7) caught no more triatomines than black traps with white LED lights. The blue traps with blue LED strips were highly attractive to other arthropods, resulting in significant reduction in triatomines caught per 1,000 other arthropods. Thus, they are less suitable for catching triatomines but might be useful in ecological studies that investigate taxonomic diversity and biomass of night-flying insects.

Hamer et al. (2024) demonstrated that the Kissing Bug Kill Trap captured four species of triatomines: *T. gerstaeckeri, T. sanguisuga, H. neotomae* and *T. rubida*. Our study adds five additional species to this list: *P. lecticularia* and *T. indictiva* in the USA, *D. maxima* and *H. peninsularis* in Mexico, and *T. dimidiata* in Guatemala. The low number (Ariano-Sánchez and Gil-Escobedo 2021) of *T. dimidiata* in Comapa, Guatemala may be due to low triatomine populations in sylvatic habitats in the region (Juarez et al. 2018) due to anthropogenic disturbance to natural habitat and the subsistence hunting of wild animals resulting in low wildlife occurrence (Mérida Ruíz et al. 2016). The eight traps in Guatemala were in continuous operation for over two years with minimal maintenance, indicating their durability and suitability for use in rural areas of Central America. Two manual searches conducted inside the Guatemala households with traps installed outside yielded zero triatomines, suggesting that in addition to serving as a more effective surveillance tool than manual searching, the traps may have intercepted dispersing triatomines before they entered households and established populations. Sampling of other households in these same Comapa, Guatemala communities in 2022 disclosed infestation rates ranging from 17–38% and colonization levels between 9–29% (Juarez et al. 2025).

Prior to development of the Kissing Bug Kill Trap, no standardized surveillance tool has existed to attempt to quantify spatio-temporal heterogeneity of adult triatomine abundance. We have now deployed large numbers of these traps over six consecutive seasons in multiple countries and have begun to capture the species diversity of dispersing adult triatomines as well as variation in adult dispersal phenology. Confirming anecdotal observations in prior years, trap catches in south Texas and San Antonio comprised mainly *T. gerstaeckeri*, with varying magnitudes and peaks (Figure 7). Within a season, the earlier occurrence in traps of *T. gerstaeckeri* in south Texas compared to San Antonio matches our expectation based on prior observations. Using traps as a standardized surveillance tool will remove observer bias associated with other forms of collection.

While we captured 1,531 triatomines during three years of field experiments, we also captured 604,432 non-target arthropods (Supplemental Table 1). While this amount of by-catch sounds large, several factors need to be considered while interpreting this study evaluating multiple trap prototypes. Among the by-catch, 43.3% were collected from the TRAP7 (blue funnel with three blue light stripes) in 2023, however, this trap captured only 10 triatomines and was excluded from further evaluation due to the excessive number of by-catch. The number of by-catch fluctuated throughout the years, especially in 2022 (Figure 8). To evaluate the phenology of triatomines in regions with poor understanding of seasonal changes in dispersal behavior, these traps we set-up and ran continuously for multiple months or multiple years. This study design therefore results in traps being deployed during periods of active triatomine dispersal and also periods of no triatomine captures (‘negative data’) but when other non-target arthropods continued to be captured. For example, large numbers of non-target arthropods were captured in the traps in August and September, 2022, in South Texas and zero triatomines were captured by those traps during those two months (Figure 7 and 8).

**Figure 8.**
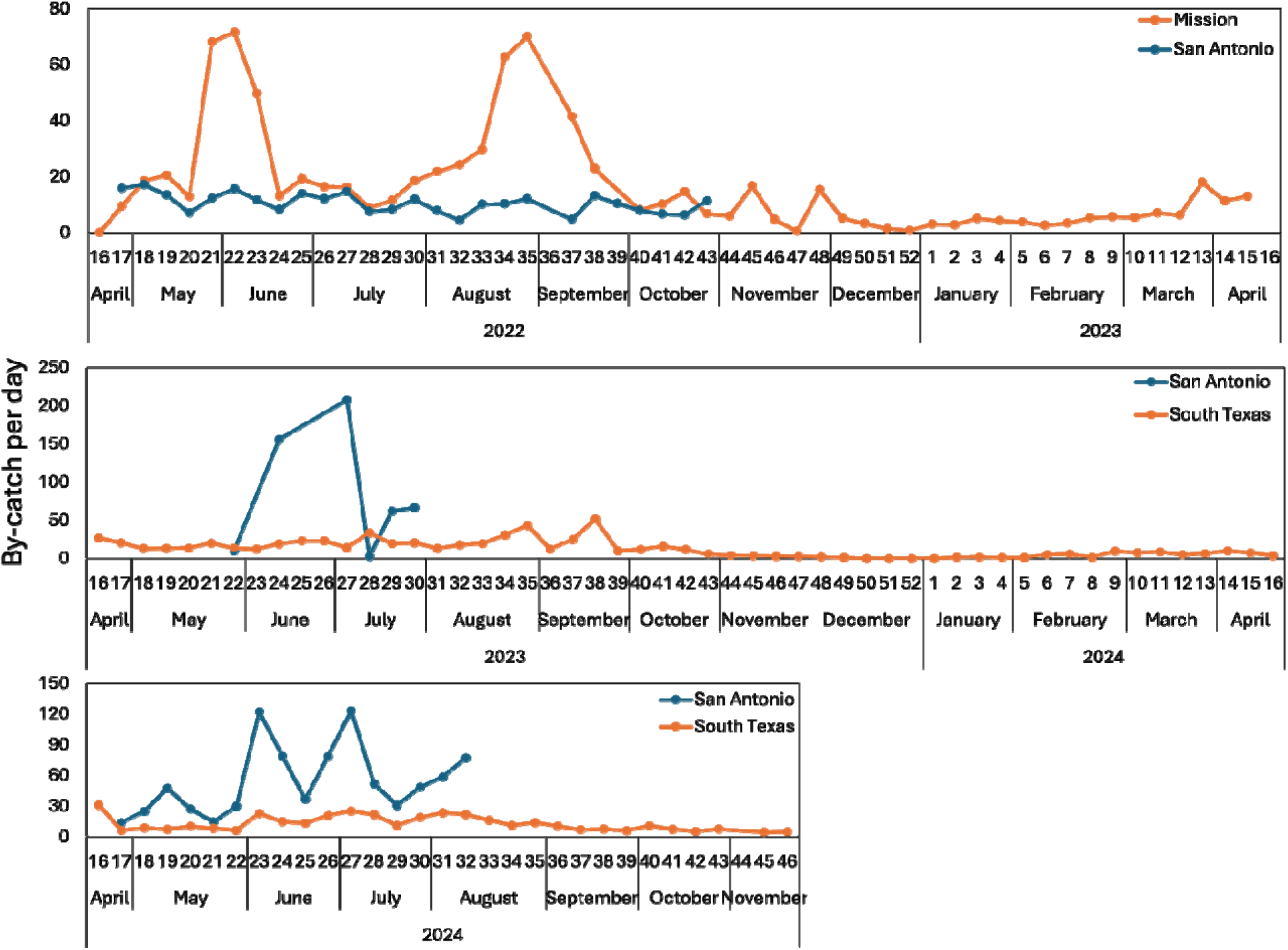
Weekly number of by-catch per day per trap in 2022-2024 at south Texas and San Antonio. The trap data for all south Texas locations and all San Antonio locations are averaged.

Even when studies quantify trap by-catch and present ratios of vectors to non-target captures (Frost 1964), this is only done during short windows of time and we are unaware of any arthropod vector trap that is continuously operated for multiple years. A vector trap that is continuously operational for multiple years is naturally going to result in a larger non-target capture. While the non-target by-catch was not needed for this study focused on triatomines, this by-catch could have alternative uses. For example, the arthropods such as Coleoptera with a hard exoskeleton are well-preserved in the propylene glycol and remain in good condition for potential contribution to an insect collection at a museum or university. Additionally, alternative preservation methods could allow these other arthropods to be repurposed as a sustainable source of protein for animal feed. Cohnstaedt et al. (2024), introduced the United States Department of Agriculture–Biomass Harvest Trap (USDA-BHT) for the mass collection of flying insects as animal feed. This innovative approach was recognized with the USDA ARSX award (United States Department of Agriculture 2022), an internal competition among USDA Agricultural Research Service employees. By-catch from these Kissing Bug Kill Traps represent a potential resource that could contribute to sustainable agriculture and waste reduction through insect biomass utilization.

It is difficult to compare the by-catch observed with these different Kissing Bug Trap prototypes because other vector trapping studies rarely quantify by-catch (but see Hribar (2020)). While trap by-catch often makes the removal of vectors time-consuming, that is not the case with triatomines in these Kissing Bug Kill Traps. Triatomines are larger than most of the arthropod by-catch (see Supplemental Figures 1-2) and also have a distinctive morphology. Therefore, when processing traps, the triatomines are easily removed as the first quick step and then the very consuming step is identifying, sorting, and counting the other arthropods (Supplemental Figure 1). Another concern with a large by-catch is the potential ecological impact by removing insects that perform essential ecological services in an environment. Fortunately, we noticed very few honeybees (1,193) in the traps over three years, so this is one beneficial insect not expected to be significantly impacted by the trap. We also contend that the impact of the traps would be inconsequential, even if the traps were used in large-scale surveillance or mass-trapping programs. The impact on arthropods from these Kissing Bug Kill Traps would represent a tiny fraction of those posed by large-scale habitat alterations, such as clearing forests for agriculture or urban development (Wagner 2020), or the impact of annual worldwide application of 3 billion kg of pesticides (Carvalho 2017) and the corresponding adverse impact on non-target arthropods (Boyce et al. 2007, Kwan et al. 2009, Pan et al. 2023, Van Deynze et al. 2024). Moreover, unlike chemical pesticides (Serrao et al. 2022, Bartling et al. 2024), the Kissing Bug Kill Trap would leave no long-term environmental residue. If the goal is to minimize trap by-catch, future prototypes of the Kissing Bug Kill Trap could have modifications to the collection cup to minimize by-catch. For example, ongoing Kissing Bug Kill Trap experiments in 2025 are using dry collection cups, which contain no liquid preservatives. If a small hole was placed in these dry collection cups, arthropods smaller than an adult triatomines, which would be the vast majority of all the by-catch, could escape unharmed while all larger arthropods remain trapped.

Of the *T. gerstaeckeri* captured, 74% were males. This result is similar to that reported for *T. infestans* and *T. eratyrusiformis* in Argentina, where more flying males were captured using light traps than females, but not for *T. guasayana*, wherein 62% were female (Abrahan et al. 2011). Fimbres-Macias et al. (2023) collected 1.6x more female than male *T. sanguisuga* manually around a building in Texas, USA, with light as an attractant. With 3,215 triatomines collected from a citizen science program, more females than males were collected for *T. gerstaeckeri*, *T. sanguisuga*, *T. indictiva*, *T. rubida*, while more males than females were collected for *P. lecticularia*, *T. protracta*, and *H. neotomae* (Curtis-Robles et al. 2018). Collectively, these results indicate that the sex ratio of triatomines attracted by light may differ among species and may differ from the sex ratio of natural populations.

Our previous study detected *T. cruzi* in triatomines captured by Kissing Bug Kill Traps (Hamer et al. 2024), demonstrating that propylene glycol is suitable for preservation of nucleic acid. The current study represents our first attempt at using bloodmeal metabarcoding (Balasubramanian et al. 2022) to identify prior hosts of triatomines captured in Kissing Bug Kill Traps. One of the two captured *T. dimidiata* subjected to the bloodmeal analysis had evidence of past feeding on dog and chicken blood, suggesting that preservation in propylene glycol is suitable for this analysis. Similarly, with a larger set of triatomines collected alive manually from houses in the same neighborhoods, Juarez et al. (2025) also found chicken and dog as frequent bloodmeal hosts, also with *Rattus rattus*, *Felis catus*, *Homo sapiens*, *Mus musculus*, *Sus scrofa*, Anatidae and *Archimandrita* sp. These bloodmeal results from the two insects caught in the Kissing Bug Kill Traps in our study suggest that they emerged from a domestic habitat, prior to flying and being intercepted by the trap, as opposed to a sylvatic habitat where wildlife hosts would be expected hosts. In the future, such assays could be used to identify the habitat of origin of these dispersing adults, which would be important in understanding not only movement of triatomines but also the origins and prevalence of *T. cruzi*.

Our results build on the initial development of the Kissing Bug Kill Trap (Hamer et al. 2024), and suggest that because of the efficacy, low cost, ease of maintenance, and durability of the trap it has promising potential for use as a large-scale surveillance tool. The trap would enable standardized passive surveillance methodology, avoiding observer bias that accompanies other active sampling techniques. The multiple funnel design and solar-powered LED lights resulted in minimal trap failure over the three year study and only one example of left of the solar-powered LED light. Most homeowners, especially in Guatemala which lacked electricity, appreciated the trap because they offered a free security light at night. The Kissing Bug Kill Trap may also have promise as a control tool to prevent dispersing triatomines from colonizing residences or domestic animal harborages thereby reducing the global threat of Chagas disease. The possibility of use as a control tool is supported by a 1954 study in which three light traps around an Arizona home captured 398 *Triatoma protracta* and the homeowners noticed fewer triatomines around their residence (Sjogren and Ryckman 1966). Deploying multiple traps around a household has proven to be effective in protecting humans from mosquito-borne pathogens (Barrera et al. 2019). The potential for practical implementation of the Kissing Bug Kill Trap awaits commercialization of this trap and its introduction into the pest control marketplace.

## Acknowledgments

We thank Katelyn Humlicek, Bisharo Farah, Emory Phu, Bryon Martinez, Sebastian Flores, Danya Garza, Orlando Lugo-Lugo, Jorge Mendoza, Javier de León, Raul Garza, Roy Rodriguez, and Phoebe Fry for deployment and monitoring of traps and processing trap contents.

## Data availability

The data used for the analyses in this study is available from the supplementary file. Sequencing data from the bloodmeal metabarcoding work along with sample and code information for analysis has been deposited in datadryad (https://datadryad.org/) with the doi: https://doi.org/10.5061/dryad.pzgmsbczg.

## Author’s contribution

Conceptualization: GLH, SAH, JHB, MGB; Data curation: YT, SS, CN; Formal analysis: YT, SB; Funding acquisition: GLH,, MGB, NP, PP; Investigation: YT, NAFS, HE, AMMV, MGP, AE, JPFM, WR, PL,TC, MK, CLH, CHD, SS, CN, SB, CA, CPC, RGR, TZS; Methodology: GLH, SAH, MGB; Visualization: YT; Writing-original draft: YT; Writing-review and editing: all authors.

