## Supplemental tables for "Advancing the design of the kissing bug kill trap for surveillance of triatomines"

Supplementary

Table 1. Arthropod bycatch for the different multi-funnel trap prototypes used in 2022 to 2024.

| Year | Trap | Triatomine | Hemiptera | Lepidoptera | Diptera | Coleoptera | Orthoptera | Odonata | Mantodea | Blattodea | Neuroptera | Apis sp. | Other Hymenoptera | Other | Total Other Arthropod | Triatomine/1000 other arthropods |
| --- | --- | --- | --- | --- | --- | --- | --- | --- | --- | --- | --- | --- | --- | --- | --- | --- |
| 2022 | TRAP1 | 125 | 710 | 1394 | 494 | 4560 | 198 | 19 | 86 | 1321 | 261 | 51 | 4294 | 698 | 14086 | 8.87 |
| 2022 | TRAP2 | 68 | 385 | 991 | 545 | 6169 | 109 | 24 | 69 | 1741 | 349 | 23 | 4166 | 527 | 15098 | 4.50 |
| 2022 | TRAP3 | 17 | 179 | 397 | 16 | 366 | 27 | 1 | 7 | 335 | 0 | 0 | 109 | 69 | 1506 | 11.29 |
| 2023 | TRAP4 | 99 | 1998 | 1225 | 2987 | 9780 | 361 | 51 | 85 | 1019 | 163 | 285 | 4753 | 3653 | 26360 | 3.76 |
| 2023 | TRAP5 | 46 | 1680 | 1824 | 1751 | 13500 | 192 | 20 | 50 | 630 | 129 | 68 | 4990 | 4564 | 29398 | 1.56 |
| 2023 | TRAP6 | 85 | 1278 | 2363 | 1938 | 9551 | 210 | 30 | 135 | 1337 | 90 | 309 | 5157 | 3347 | 25745 | 3.30 |
| 2023 | TRAP7 | 10 | 15389 | 4018 | 4682 | 122759 | 219 | 38 | 0 | 642 | 592 | 3 | 47648 | 65505 | 261495 | 0.04 |
| 2023 | TRAP2A | 252 | 3521 | 3641 | 2938 | 19665 | 458 | 88 | 171 | 2411 | 249 | 135 | 10411 | 4285 | 47973 | 5.25 |
| 2024 | TRAP10 | 86 | 1298 | 983 | 899 | 7808 | 381 | 23 | 182 | 1203 | 647 | 114 | 4178 | 1186 | 18902 | 4.55 |
| 2024 | TRAP8 | 60 | 1968 | 1305 | 1487 | 9067 | 199 | 26 | 869 | 1096 | 173 | 29 | 6824 | 1248 | 24291 | 2.47 |
| 2024 | TRAP2A | 89 | 1596 | 1355 | 2418 | 8325 | 227 | 39 | 449 | 1138 | 341 | 50 | 6320 | 1640 | 23898 | 3.72 |
| 2024 | TRAP9 | 98 | 1136 | 1024 | 1061 | 6925 | 145 | 39 | 113 | 880 | 581 | 18 | 4552 | 596 | 17070 | 5.74 |
| 2024 | TRAP11 | 32 | 1768 | 426 | 555 | 5246 | 109 | 6 | 140 | 441 | 85 | 42 | 1714 | 906 | 11438 | 2.80 |
| 2024 | TRAP2A | 149 | 1677 | 1529 | 1284 | 10283 | 237 | 29 | 255 | 771 | 233 | 28 | 3536 | 913 | 20775 | 7.17 |
| 2024 | TRAP12 | 71 | 4437 | 1906 | 2732 | 12463 | 250 | 52 | 846 | 1250 | 231 | 21 | 6308 | 2100 | 32596 | 2.18 |
| 2024 | TRAP13 | 105 | 2975 | 2161 | 3079 | 14012 | 318 | 42 | 461 | 921 | 272 | 17 | 7385 | 2158 | 33801 | 3.11 |

Table 2. The number of triatomines and triatomines per day per trap (in parenthesis) of the six species captured from 2022-2024 in Texas and Guatemala.

|  | Jan | Feb | Mar | April | May | June | July | Aug | Sep | Oct | Nov | Dec |
| --- | --- | --- | --- | --- | --- | --- | --- | --- | --- | --- | --- | --- |
| 2022 |  |  |  |  |  |  |  |  |  |  |  |  |
| *Triatoma dimidiata* | - | - | - | - | - | - | - | - | - | 1  (0.0040) | 1  (0.0042) | - |
| *Triatoma gerstaeckeri* | - | - | 2 (0.0036) | 5 (0.0167) | 41 (0.1323) | 83 (0.2767) | 16 (0.0516) | - | - | - | - | - |
| *Triatoma indictiva* | - | - | - | - | - | - | - | - | - | - | - | - |
| *Paratriatoma lecticularia* | - | - | - | - | - | - | - | - | - | - | - | - |
| *Hospesneotomae neotomae* | - | - | - | - | - | - | - | - | - | - | - | - |
| *Triatoma sanguisuga* | - | - | - | - | - | - | 3  (0.097) | - | - | - | - | - |
| 2023 |  |  |  |  |  |  |  |  |  |  |  | - |
| *Triatoma dimidiata* | - | - | 1  (0.0040) | 2  (0.0083) | - | - | 1  (0.0040) | 1  (0.0040) | - | - | - | - |
| *Triatoma gerstaeckeri* | - | - | 2  (0.0015) | 146  (0.1106) | 48  (0.0352) | 125  (0.0947) | 77  (0.0565) | 24  (0.0176) | 1  (0.0008) | - | - | - |
| *Triatoma indictiva* | - | - | - | - | - | - | - | - | - | - | - | - |
| *Paratriatoma lecticularia* | - | - | - | 2  (0.0015) | 1  (0.0007) | 11  (0.0083) | - | 1  (0.0007) | 1  (0.0008) | - | - | - |
| *Hospesneotomae neotomae* | - | - | - | 2  (0.0015) | - | 3  (0.0023) | 1  (0.0007) | 1  (0.0007) | - | - | - | - |
| *Triatoma sanguisuga* | - | - | - | - | - | - | 4  (0.0029) | 2  (0.0015) | - | - | - | - |
| 2024 |  |  |  |  |  |  |  |  |  |  |  |  |
| *Triatoma dimidiata* | 1  (0.0040) | 1  (0.0043) | - | 2  (0.0081) | - | - | - | - | - | - | 2  (0.0083) | - |
| *Triatoma gerstaeckeri* | - | - | - | 14  (0.0083) | 110  (0.0634) | 262  (0.1560) | 153  (0.0881) | 43  (0.0248) | 2  (0.0012) | 2  (0.0012) | - | - |
| *Triatoma indictiva* | - | - | - | - | - | - | 3  (0.0017) | 3  (0.0017) | - | - | - | - |
| *Paratriatoma lecticularia* | - | - | - | 2  (0.0012) | 5  (0.0029) | 6  (0.0036) | 1  (0.0006) | 2  (0.0012) | - | - | - | - |
| *Hospesneotomae neotomae* | - | - | - | - | 1  (0.0006) | - | 2  (0.0012) | 1  (0.0006) | 1  (0.0006) | 1  (0.0006) | - | - |
| *Triatoma sanguisuga* | - | - | - | - | 1  (0.0006) | 2  (0.0012) | 1  (0.0006) | 3  (0.0017) | - | - | - | - |

Triatomine per day per trap for the month = Triatomine number / days in the month / number of traps in the year

Number of trap in Texas: 10 (2022), 44 (2023), 56 (2024)

*Triatoma dimidiata* was calculated using the number of traps deployed in Guatemala (n = 8)

- Indicates no triatomine were captured in the month

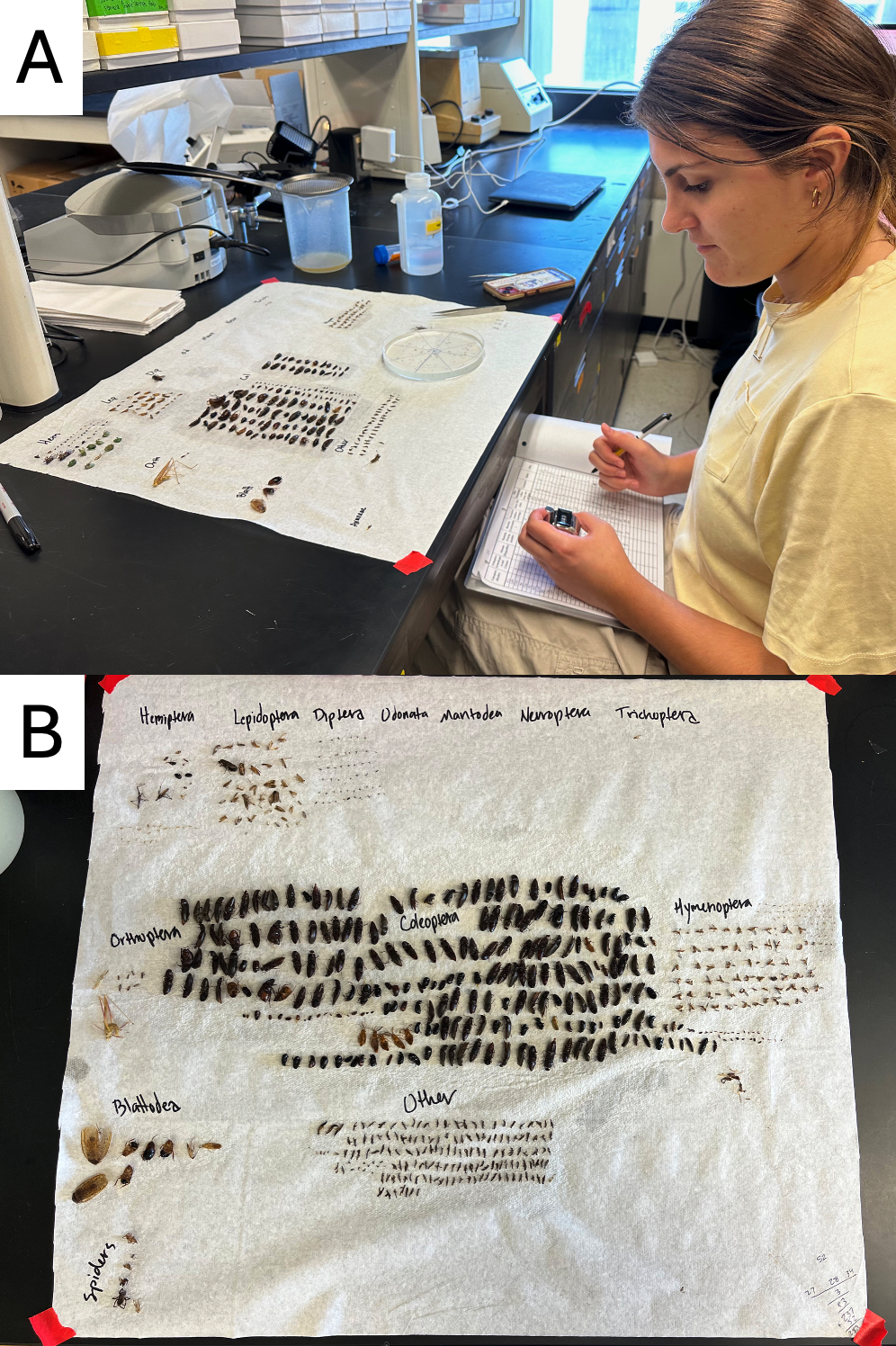

Supplemental Figure 1. Examples of processing by-catch arthropods from Kissing Bug Kill Traps after a week of collection. Triatomines were removed and counted as the first step as they are larger than most of the by-catch and have a distinctive shape.

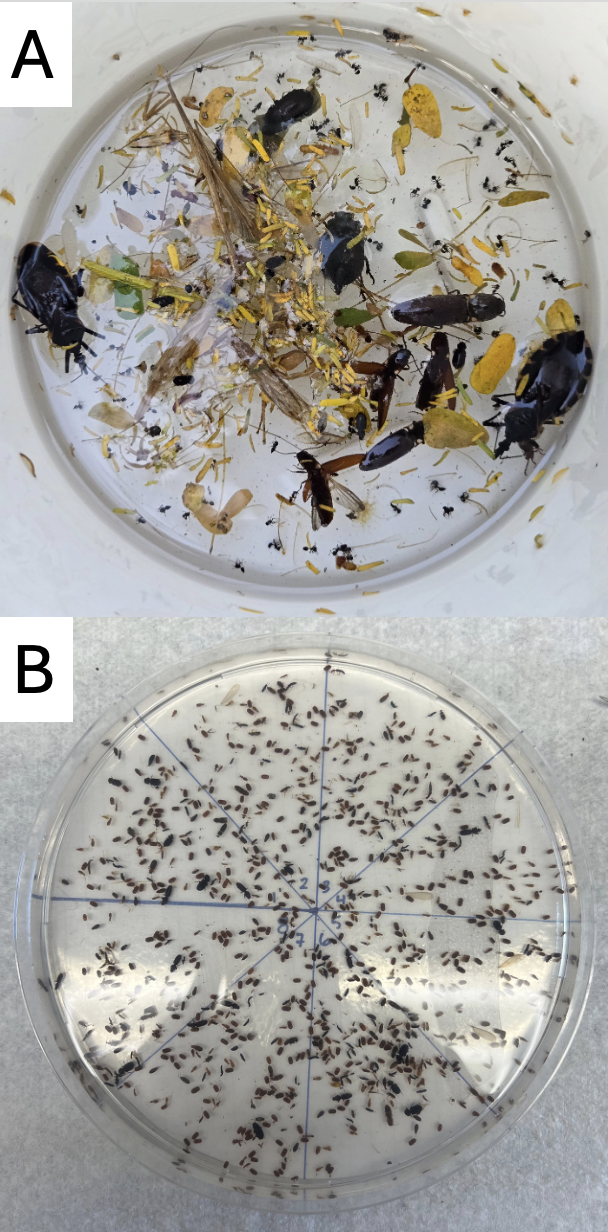

Supplemental Figure 2. Example of the collected arthropods, including two adult triatomines, from a Kissing Bug Kill Trap deployed for one week (A) and an example of the floating method to speed-up processing of large numbers of small arthropod by-catch (B).
